# Impact of dose, duration and host immune status on ultrashort telacebec treatment in a mouse model of Buruli ulcer

**DOI:** 10.1101/2021.06.22.449541

**Authors:** Deepak V. Almeida, Paul J. Converse, Oliver Komm, Eric L. Nuermberger

**Affiliations:** Center for Tuberculosis Research, Department of Medicine, Johns Hopkins University, Baltimore, Maryland, USA; Department of Tropical Medicine, Bernhard Nocht Institute for Tropical Medicine & I. Department of Medicine, University Medical Center Hamburg-Eppendorf, Hamburg, Germany

## Abstract

Telacebec (Q203) is a new anti-tuberculosis drug in clinical development with extremely potent activity against *Mycobacterium ulcerans*, the causative agent of Buruli ulcer (BU). The potency of Q203 has prompted investigation of its potential role in ultra-short, even single-dose, treatment regimens for BU in mouse models. However, the relationships of Q203 dose and duration and host immune status to treatment outcomes remain unclear, as does the risk of emergence of drug resistance with Q203 monotherapy. In the present study, immunocompetent BALB/c and immunocompromised SCID-beige mice were infected in both hind footpads and treated eight weeks later. In both mouse strains, controls received rifampin-clarithromycin; others received Q203 at 0.5 or 2 mg/kg/d for 5 or 10 days. Additionally, BALB/c mice received a single dose of 2.5 or 10 mg/kg or 3.3 mg/kg/d for 3 days. Treatment response was based on changes in footpad swelling and CFU counts at the end of treatment as well as 4 and 13 weeks after stopping treatment. Efficacy depended on total dose more than duration. Total doses of 5-20 mg/kg rendered nearly all BALB/c mice culture-negative 13 weeks post-treatment without selection of Q203-resistant bacteria. Although less potent in SCID-beige mice, Q203 still rendered the majority of footpads culture-negative at total doses of 10-20 mg/kg. However, Q203 resistance was identified in relapse isolates from some SCID-beige mice. Overall, these results support the potential of Q203 monotherapy for single-dose or other ultra-short therapy for BU, although highly immunocompromised hosts may require higher doses or durations and/or combination therapy.

## INTRODUCTION

Treatment of Buruli ulcer (BU), a disease primarily endemic to regions of sub-Saharan Africa and parts of Australia (1), has evolved from extensive surgical excision to the first effective combination chemotherapy regimen of rifampin (RIF, R) and streptomycin given daily for 8 weeks to the currently recommended oral regimen of RIF and clarithromycin (CLR, C) for 8 weeks (2, 3). Though clinical studies have shown good efficacy of the RIF+CLR regimen (4), shortening the duration of treatment and reducing the potential for adverse effects and drug-drug interactions would make it easier to implement.

Telacebec (Q203, Q), a new drug in clinical development to treat tuberculosis (5), is extremely potent against *Mycobacterium ulcerans*, the causative agent of BU, due to the high vulnerability of the bacterial respiratory cytochrome bc_1_:aa_3_ oxidase to this agent (5) in the absence of a functional alternate terminal oxidase (6). In our previous experiments we demonstrated the efficacy of Q203 in a mouse footpad infection model of BU, including its potential to drastically reduce the treatment duration needed for cure (7, 8). Q203 at doses as low as 0.5 mg/kg given alone for 5 days showed remarkable bactericidal activity, extending well beyond the last day of drug ingestion (7). This, coupled with reports from other investigators has led to suggestions to explore the possibility of a simple one-dose treatment of BU (7, 9, 10). However, it is not clear if the same total dose of Q203 is as effective when given as a single dose compared to daily divided doses; secondly, the optimal equipotent dose when compared to humans is not known; and, finally, it is not clear whether Q203 monotherapy would lead to selection of drug resistance and result in treatment failure.

We designed the current study in the mouse footpad infection model to answer these questions. We performed a dose-ranging and dose-fractionating study of Q203 to determine the total dose necessary for cure and whether a single dose or divided daily doses is a more effective way to deliver the same total dose. To evaluate the ability of Q203 monotherapy to sterilize infected footpads without selecting drug-resistant mutants in mice with deficient innate and adaptive immune responses, we included SCID-beige mice in selected treatment groups. In our previous study, mice were observed for a maximum of 4 weeks post-treatment. In the present study, we extended the observation period to 13 weeks post-treatment to give a longer time for mice to relapse if viable bacteria remained in the footpads after treatment.

Our results demonstrate that single-dose treatment may indeed be as effective as daily divided doses and successfully resolve BU. However, Q203 was less effective and selected for drug-resistant mutants in SCID-beige mice, indicating that monotherapy may not be appropriate for use in highly immunocompromised hosts.

## RESULTS

### Footpad swelling and CFU counts in BALB/c mice

The mean (±SD) CFU count on the day after footpad infection was 3.97 ± 0.52 log_10_ per footpad. Mice exhibiting footpad swelling grades between 2 and 3 at 8 weeks post-infection were included in the study, and randomized to receive either the RC control regimen or one of the test regimens (Table S1). Four total doses were tested: 2.5, 5, 10 and 20 mg/kg. The 2.5 mg/kg total dose was given as a single dose or divided into five consecutive daily doses of 0.5 mg/kg. The 5 mg/kg total dose was given as ten daily (5/7) doses of 0.5 mg/kg over 2 weeks. The 10 mg/kg total dose was given as a single dose or divided into three consecutive daily doses of 3.3 mg/kg or five daily doses of 2 mg/kg. The 20 mg/kg total dose was given as ten daily (5/7) doses of 2 mg/kg over 2 weeks. At the start of treatment (D0), the median swelling grade was 2.5. Q203 rapidly reduced swelling in all dosing groups to a median swelling grade of ≤ 1 within the first week of treatment and to a negligible swelling grade by Weeks 3-5 irrespective of whether treatment was ≤ 1 week (Fig 1A) or 2 weeks (Fig 1B) in duration. Mice receiving RIF+CLR experienced a slower but persistent improvement in swelling after completing 1 or 2 weeks of treatment; and those completing 2 weeks reached a state of negligible swelling between Weeks 6-8, but the median increased back to 0.5 by the relapse time point at Week 14, as it did for the group that received a single 2.5 mg/kg dose of Q203.

**Fig 1.**
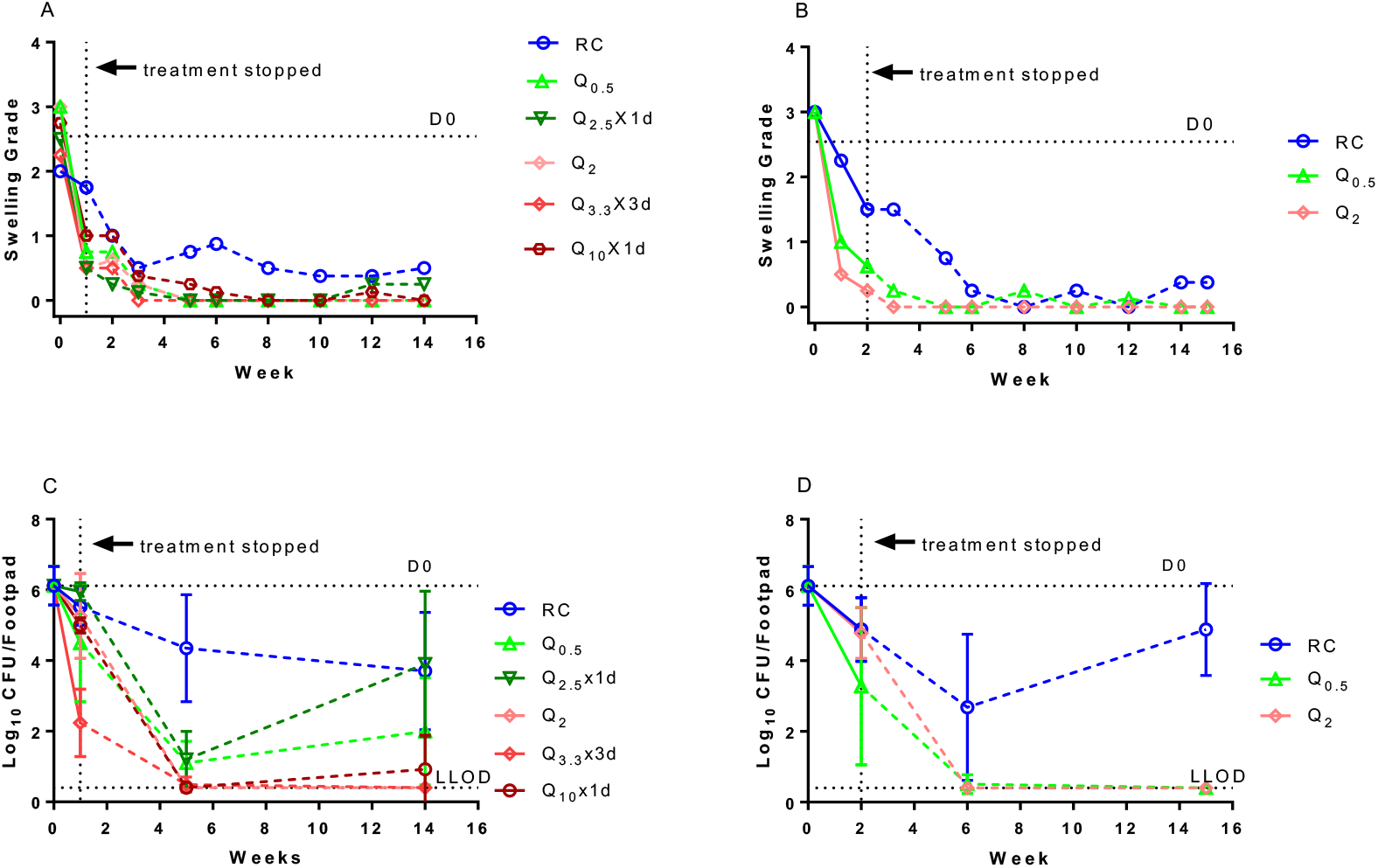
Footpad swelling grade and microbiological outcome in BALB/c mice in response to treatment. Median swelling grade in BALB/c mice treated for 1 week (A) and BALB/c mice treated for 2 weeks (B). Log_10_ CFU/footpad in BALB/c mice treated for 1 week (C) and BALB/c mice treated for 2 weeks (D). Solid lines represent change during treatment, while that after stopping treatment is shown by dashed lines. RC, RIF 10 mg/kg plus CLR 100 mg/kg. Numbers in subscript after the drug abbreviation are the doses in mg/kg. D0, day 0 or the beginning of treatment. The lower limit of detection (LLOD) at Week 1 was 1.48 log_10_ CFU. At Week 2, it was 0.48 log_10_ CFU. For other time points, it was 0.40 log_10_ CFU.

The mean CFU count at D0 was 6.12 ± 0.55 log_10_ per footpad (Figs 1C & D). Mean CFU counts in RIF+CLR-treated mice declined in a duration-dependent manner on treatment and over the next 4 weeks post-treatment before reaching a plateau or increasing again at the relapse time point at 13 weeks post-treatment. Two out of 6 footpads were culture-negative 4 weeks after completing 2 weeks of RIF+CLR treatment, but all 8 footpads were positive at 13 weeks post-treatment with a mean CFU count of nearly 5 log_10_ (Table S2). As suggested by the swelling results, Q203 was more effective. Just 1 week (i.e., 5 doses) of Q_0.5_ (henceforth the dose in mg/kg is shown in subscripts) rendered 2 of 6 and 2 of 8 footpads culture-negative at 4 and 13 weeks post-treatment, respectively. Extending this dose for another week rendered all 8 footpads culture-negative 13 weeks later, except for 1 footpad with 1 CFU. Overall, 5 doses of Q_0.5_ was significantly more effective than 10 doses of RIF+CLR when assessed at Week 1+4 (p<0.0001), but not at the relapse time point (Week 1+13). Ten doses of Q_0.5_ was significantly more effective than 10 doses of RIF+CLR when assessed at Week 2+4 and at the relapse time point (2+13) (p=0.015 and p<0.001, respectively). The higher dose of Q_2_ daily for 1 or 2 weeks rendered all footpads culture-negative at 4 weeks post-treatment and prevented relapse at 13 weeks post-treatment, except for a single footpad with 1 CFU at Week 1+13. While superior to RIF+CLR (p<0.01 at Weeks 1+4, 1+13, 2+4, 2+13), Q_2_ was not statistically superior to Q_0.5_ at these time points. A single dose of Q_2.5_ resulted in a higher mean CFU/footpad count at Week 1+13 than the same total dose divided over 5 days, but the difference was not statistically significant. A single dose of Q_10_ or 3 daily doses of Q_3.3_ appeared to be just as effective as the same total dose divided over 5 days (Q_2_). Each of these regimens resulting in a total dose of 10 mg/kg was superior to RIF+CLR at Weeks 1+4 (p<0.0001) and 1+13 (p<0.05) and to the single 2.5 mg/kg dose at Week 1+13 (p<0.05).

### Footpad swelling and CFU counts in SCID-beige mice

SCID-beige mice lacking adaptive T-cell and B-cell responses and functional natural killer cells were included in selected treatment groups. The mean CFU count the day after footpad infection was 4.17 ± 0.22 log_10_ CFU/footpad. At D0, the median swelling grade was 2.6 (Figs 2A and B) and mice were randomized to receive either RIF+CLR, Q_0.5_ or Q_2_ for up to two weeks. Single-dose regimens were not included for testing in SCID-beige mice. As observed in BALB/c mice, both Q203 doses rapidly reduced the swelling grade, which became negligible within 1-2 weeks of completing 1-2 weeks of treatment with either dose and remained negligible through to the relapse time point, Week 14, except in the group receiving Q_0.5_ for 1 week, which uniformly experienced recrudescence of swelling at Week 10 and required euthanasia thereafter. RIF+CLR reduced the swelling grade less rapidly than Q203 and only while mice were on treatment, as footpad swelling rapidly rebounded within 1-3 weeks of stopping RIF+CLR treatment and eventually necessitated euthanasia.

**Fig 2.**
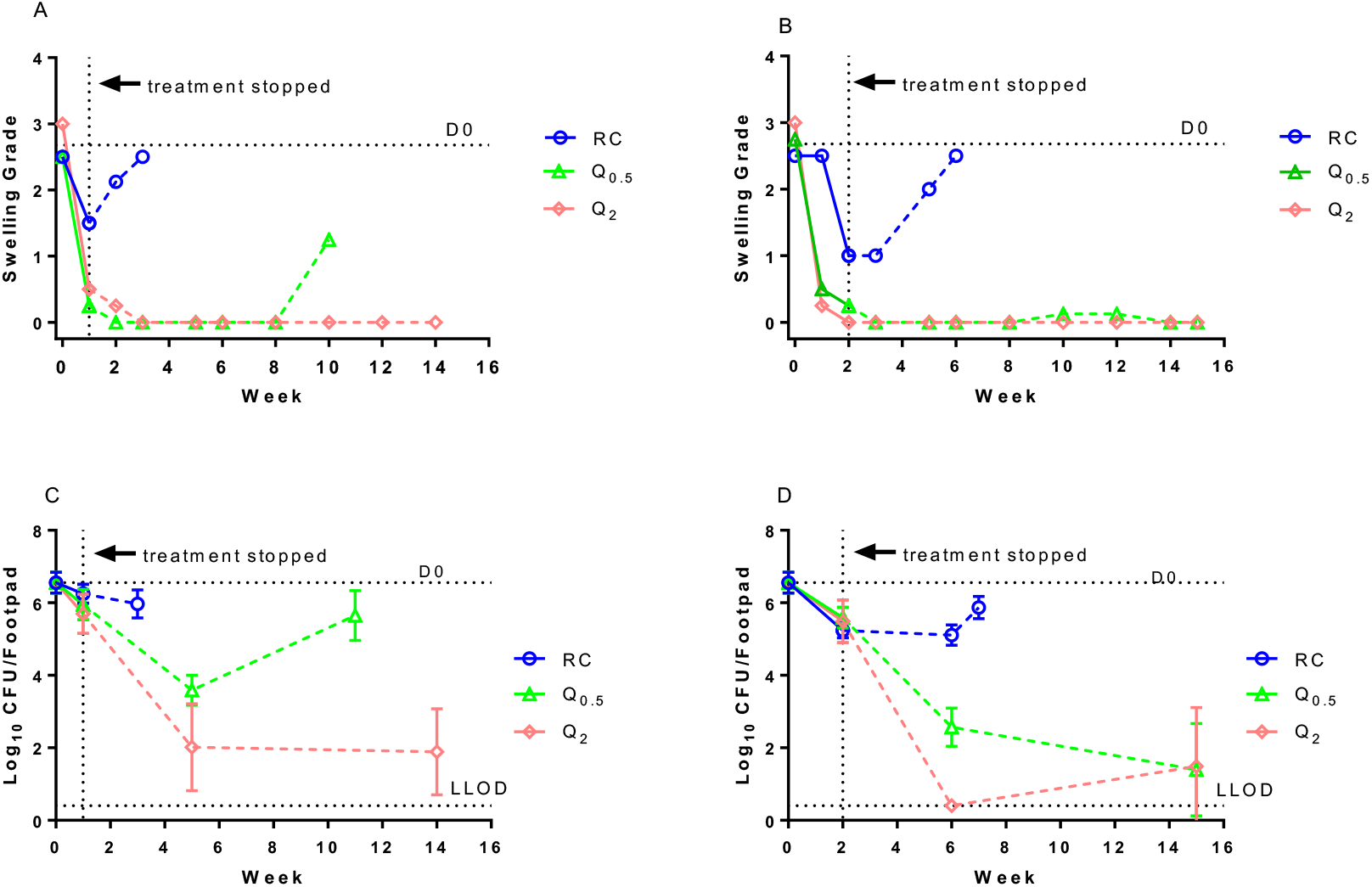
Footpad swelling grade and microbiological outcome in SCID-beige mice in response to treatment. Median swelling grade in SCID-beige mice treated for 1 week (A) and SCID-beige mice treated for 2 weeks (B). Log_10_ CFU/footpad in SCID-beige mice treated for 1 week (C) and SCID-beige mice treated for 2 weeks (D). Solid lines represent change during treatment, while that after stopping treatment is shown by dashed lines. RC, RIF 10 mg/kg plus CLR. Numbers in subscript after the drug abbreviation are the doses in mg/kg. D0, day 0 or the beginning of treatment. Lower limit of detection (LLOD) at W1 was 1.48 log_10_ CFU, at W2 was 0.48 log_10_ CFU, and for other time points, it was 0.40 log_10_ CFU.

The mean CFU count at D0 was 6.56 ± 0.29 log_10_ per footpad (Figs 2C and D). RIF+CLR treatment resulted in duration-dependent reductions in footpad CFU counts of up to 1.3 log_10_ after 2 weeks, but the CFU counts increased again to approximately 6 log_10_ by the time the humane endpoint was reached at Week 2+5. Q_0.5_ for 1 week resulted in a greater bactericidal effect (i.e., 3.0 log_10_ reduction) at Week 1+4 but the CFU counts were again approaching 6 log_10_ when euthanasia was required at Week 1+9. Increasing the total dose by extending the duration of Q_0.5_ to 2 weeks or giving Q_2_ for 1 or 2 weeks prevented such treatment failures and reduced the CFU counts at 4 weeks post-treatment in a dose- and duration-dependent fashion. Unlike in BALB/c mice, Q_2_ was superior to Q_0.5_ at both Week 1+4 and Week 2+4 (p<0.05 and p<0.0001, respectively) in SCID-beige mice. At the relapse time point, 13 weeks post-treatment, between 4 and 5 of the 8 footpads in each group were culture-negative, suggesting that the pathogen had been cleared from at least half of the mice.

Despite its powerful bactericidal and sterilizing activity in both mouse strains, Q203 was less potent in SCID-beige mice. BALB/c mice had significantly lower mean CFU counts than SCID-beige mice after treatment with Q_0.5_ (p<0.05 at Weeks 1 and 2; p<0.001 at Weeks 1+4 and 2+4) and Q_2_ (p=0.001 at Week 1+4).

### Resistance detection

By plating serial 10-fold dilutions of a culture suspension of *M. ulcerans* on agar plates supplemented with Q203 at 4 and 40 times the previously determined MIC of 0.075 ng/ml (8), the frequency of spontaneous Q203-resistant mutants *in vitro* was determined to be 1.2 in 10^5^ on plates containing Q203 at 0.3 ng/ml (i.e., 11 CFU per 900,000 total CFU plated). No CFU were detected on plates containing Q203 at 3 ng/ml, indicating that the frequency of resistance at 40x MIC was less than 1 in 10^6^. In a second experiment with a higher bacterial inoculum (6 × 10^7^ CFU/ml), the frequency of resistance to 3 ng/ml was 5 in 10^8^.

Isolates from 4 BALB/c mouse footpads obtained at the relapse time point, 13 weeks post-treatment, from mice that received 5 doses of Q_2_ (n=1), a single dose of Q_10_ (n=2), or 10 doses of Q_0.5_ were plated in serial dilutions on agar containing 0.3 and 3 ng/ml of Q203, and no Q resistant-mutants were detected. Isolates from 2 SCID-beige mice among those receiving 5 doses of Q_0.5_ that uniformly experienced early footpad re-swelling had approximately 1 in 10^5^ CFU growing at 3 ng/ml and 1 in 10^4^ CFU growing at 0.3 ng/ml, indicating that, although some selective amplification of resistant mutants may have occurred with treatment, it was not responsible for the treatment failure. After extending treatment with Q_0.5_ to 2 weeks, 4 of 8 footpads remained culture-positive and no selective amplification of resistant mutants was detected on either 0.3 or 3 ng/ml. However, treatment with Q_2_ for 2 weeks did select Q203-resistant mutants in some SCID-beige mice. At the relapse time point, 13 weeks post-infection, 3 of 8 footpads were culture-positive, and all 3 isolates were resistant to Q203 at both 0.3 and 3 ng/ml. Three colonies from each of these 3 isolates were subjected to PCR amplification and sequencing of the *qcrB* gene. All isolates had the same mutation, a single nucleotide polymorphism (A → G) at nt967 resulting in amino acid change T323A.

## DISCUSSION

Q203 is a new drug targeting bacterial respiration that is in clinical development for tuberculosis and has also proven to be very effective against *M. ulcerans* in pre-clinical studies (7-11). Previously, we showed its potential for reducing the duration of treatment to 1-2 weeks based in large part on the extended duration of bactericidal activity after stopping treatment (7). At least a portion of this extended activity of Q203 is attributable to its potent activity and slow clearance from plasma and site of infection. However, we also questioned whether treatment with Q203 might promote bacterial clearance by host-mediated immune mechanisms. An effective host response may also have an important role in preventing the selection of drug-resistant bacteria, especially if Q203 monotherapy were to be considered. In the current study, we explored the possibility of a single-dose or other ultrashort treatment duration for *M. ulcerans* compared to 1-2 weeks of treatment and assessed the impact of dose, dosing schedule and the host immune response on the efficacy of such regimens by comparing long-term post-treatment outcomes in immune-competent BALB/c mice and immune-compromised SCID-beige mice. This also afforded the opportunity to assess the potential for emergence of drug resistance during monotherapy with Q203 and the impact of host immunity.

Q203 again exhibited strong bactericidal effects that extended well beyond the end of the dosing period, even if that dosing period consisted of a single dose. In BALB/c mice, footpad swelling resolved almost completely in as little as 1-2 weeks after treatment, including in groups treated with single-dose regimens. Q203 showed clear dose-ranging activity with higher doses showing increased activity. Total doses of 5-20 mg/kg achieved relapse-free cure in the vast majority of footpads at 13 weeks post-treatment, while total doses of 2.5 mg/kg were less effective. A single dose of 10 mg/kg was equally effective as 2 mg/kg given daily for 5 days or 3.3 mg/kg for 3 days. While our study was underway, Thomas et al (9) reported that a single Q203 dose of 20 mg/kg given once or divided into four doses of 5 mg/kg given over a week resulted in relapse-free cure in a mouse footpad model. Our study supports and extends those results by showing that a lower single-dose regimen of Q at 10 mg/kg or three divided daily doses of 3.3 mg/kg was also highly effective. Our prior PK data (7) suggest that Q203 doses between 2-10 mg/kg likely correspond well to human doses of 100-300 mg that were recently reported to be well tolerated and safe in phase 1 trials and in TB patients over 14 days of dosing in a recent phase 2a trial (12). We also tested a total dose of 20 mg/kg given over a period of 2 weeks given as 2 mg/kg daily, 5 days per week, and it was the only regimen that eliminated all cultivable bacilli in all BALB/c mouse footpads (although 10 mg/kg divided over 5 days left only a single culture-positive footpad with a single detectable CFU). Taken together, these results provide strong support for the prospect of single-dose or other ultra-short therapy of *M. ulcerans* infection with doses that have thus far been safe and well tolerated in humans.

Comparisons of regimen efficacy between BALB/c and SCID-beige mice demonstrated important effects of the host immune response on the response to treatment with both Q203 and the RIF+CLR control regimen. BALB/c mice treated with RIF+CLR for 1 or 2 weeks experienced a gradual reduction in footpad swelling and CFU counts for at least 4 weeks after stopping treatment and did not have recrudescent swelling of the footpad despite the vast majority of the footpads remaining culture-positive with >1,000 CFU at 13 weeks post-treatment. In stark contrast, SCID-beige mice experienced a rebound in footpad swelling almost immediately after stopping treatment as well as little additional decline in footpad CFU counts in the first few weeks post-treatment before the burden increased again with longer incubation (i.e., in the group treated for 2 weeks). These results indicate that the magnitude of the bactericidal effect of RIF+CLR is, to a significant extent, dependent on effective natural killer cells and/or adaptive host immunity, as is the post-treatment resolution of footpad swelling and containment of bacterial growth. These immune effects may be enhanced by suppression of mycolactone synthesis by drug treatment.

The efficacy of Q203 was also impacted by mouse strain. Although resolution of swelling during Q203 treatment was at least as rapid in SCID-beige mice compared to BALB/c mice, the magnitude of its bactericidal effect in SCID-beige mice, while substantial, was not as great as that observed in BALB/c mice. Nevertheless, sustained bactericidal effects were observed after Q203 treatment in SCID-beige mice and the majority of those treated with Q203 at 2 mg/kg for 1-2 weeks were culture-negative 13 weeks post-treatment, despite having mean CFU counts of 5.5-5.7 log_10_ at the end of dosing. This striking result affirms the persistent bactericidal effects and sterilizing efficacy of Q203, even in the absence of an effective adaptive host immune response.

Another important conclusion drawn by comparing Q203 treatment responses in BALB/c and SCID-beige mice is that the host immune response plays a key role in preventing the emergence of Q203 resistance during treatment. Monotherapy of active mycobacterial infections is generally strongly discouraged because of the propensity for selecting spontaneous drug-resistant mutants. However, as we and others have argued previously (7, 9, 11), Q203 monotherapy of BU may be considered given the low spontaneous frequency of resistance mutations, their potential fitness cost, the potential for adaptive host immunity to contain or clear small residual populations of drug-resistant bacteria and the small-to-absent risk of person-to-person transmission of a drug-resistant infection, even if it occurs. As we hoped, we observed no evidence of Q203 resistance among the isolates obtained from the last few culture-positive BALB/c mice at 13 weeks post-treatment. In contrast, all SCID-beige mice remaining culture-positive after 2 weeks of treatment with Q203 at 2 mg/kg harbored Q203-resistant isolates with a mutation in *qcrB*. This resistance mutation was previously reported by Scherr et al to result in a 230.5-fold increase in the MIC of Q203 (11), which is consistent with growth of our isolates on agar containing Q203 at 40 times the MIC. The previously reported frequency of spontaneous resistant mutants was previously reported to be 1 in 10^9^ at 10 nM (5.57 ng/ml) (11). We found it to be around 1 in 10^5^ at 0.3 ng/ml and 5 in 10^8^ at 3 ng/ml in our *in vitro* selection studies. As the CFU counts at the start of treatment in SCID-beige mice were 6.56 ± 0.29 log_10_ CFU per footpad, it is possible that some of the mice harbored spontaneous Q203-resistant mutants at the start of treatment and these were selectively amplified at a higher dose of Q at 2 mg/kg under its strong selective pressure in the absence of a host immune response resulting in treatment failure in these mice. Our results in SCID-beige mice therefore indicate the possibility that Q203 monotherapy in highly immune-compromised hosts could lead to selection of drug resistance and multi-drug therapy should be considered in this instance.

## MATERIALS AND METHODS

### Bacterial strain

*M. ulcerans* strain 1059, originally obtained from a patient in Ghana, was used for the study (13).

### Antibiotics

RIF powder was purchased from Sigma. CLR pills were purchased from the Johns Hopkins Hospital pharmacy. Q203 was kindly provided by the Global Alliance for TB Drug Development. RIF and CLR were prepared separately in sterile 0.05% (wt/vol) agarose solution in distilled water. Q203 was formulated in 20% (wt/wt) D-α tocopheryl polyethylene glycol 1000 (Sigma) succinate solution (7).

### Mouse infection

To prepare the inoculum, mouse footpads infected with *M. ulcerans* 1059 were harvested upon reaching a swelling grade between 2 and 3. After thorough disinfection with 70% alcohol swabs, the footpad tissue was dissected away from bone and then homogenized by fine mincing before being suspended in sterile phosphate-buffered saline (PBS). The suspensions were then frozen in 1.5 ml aliquots and stored at −80° C. Prior to infection, vials were thawed to room temperature and pooled together to obtain the required amount for mouse infection. BALB/c and Fox Chase SCID Beige mice (Charles River Laboratories) were inoculated subcutaneously in both hind footpads with 0.03 ml of a culture suspension containing *M. ulcerans* 1059. Treatment began approximately 8 weeks (D0) after infection when the mice had a footpad swelling grade of 2-3.

### Treatment

Drugs were administered orally in 0.2 ml by gavage. Drug doses were chosen to produce similar area under the plasma concentration-time curves over 24 hours post-dose compared to human doses, as previously described (7, 8, 14). All animal procedures were conducted according to relevant national and international guidelines and approved by the Johns Hopkins University Animal Care and Use Committee. Mice were randomized to treatment groups (Table S2). The control regimen consisted of RIF 10 mg/kg plus CLR 100 mg/kg. Test regimens consisted of either Q_0.5_ or Q_2_ given 5 days per week (5/7) for 1-2 weeks, Q_3.3_ given daily for 3 consecutive days, or Q_2.5_ or Q_10_ given as a single dose. The single dose of Q_2.5_ was selected to match the total dose of Q_0.5_ given for 1 week, the Q_10_ single dose matched the total dose of Q_2_ given for 1 week and Q_3.3_ given for 3 days. SCID-beige mice were included to inform on the sterilizing potential of the regimen when compared to the activity in BALB/c mice and the risk of selection of drug resistance during monotherapy in an immunocompromised host.

### Evaluation of treatment response

Treatment outcomes were evaluated based on (i) decrease in footpad swelling, denoted as swelling grade, and (ii) decrease in CFU counts. The swelling grade was scored as described previously (14). Briefly, the presence and the degree of inflammatory swelling of the infected footpad were assessed weekly and scored from 0 (no swelling) to 4 (inflammatory swelling extending to the entire limb) for all surviving mice. For CFU counts, six footpads (from three mice) were evaluated on the day after infection, and at the start of treatment (D0) to determine the infectious dose and the pretreatment CFU counts, respectively. The response to treatment was determined by plating 6 footpads (from 3 mice) at the end of 1 week or 2 weeks treatment. Mice treated with single dose or the 3-day regimen were sacrificed at the end of 1 week along with 1 week treated mice. As shown previously (7), Q203 exhibits extended activity after stopping treatment. To evaluate this, mice from 1-week and 2-week treatment groups were held for an additional 4 weeks (3 mice) or 13 weeks (4 mice) to assess relapse without treatment. During this period footpads were inspected every 2 weeks for any signs of re-swelling after stopping treatment. When re-swelling was observed, mice were sacrificed when the swelling reached a lesion index ≥ 3 and the footpads were harvested and plated for CFU counts. All the remaining mice were sacrificed at the end of their 4- or 13-week observation period and their footpads were harvested and plated for CFU. Footpad tissue was harvested after thorough disinfection with 70% alcohol swabs and then homogenized by fine mincing before suspending in sterile phosphate buffered saline (PBS). Ten-fold serial dilutions and undiluted fractions of homogenate were plated in 0.5 ml aliquots on selective 7H11 agar supplemented with 10% OADC and incubated at 32°C for up to 12 weeks before CFU were enumerated.

### Resistance testing

The proportion of resistant mutants growing on 0.3 ng/ml (4xMIC) and 3 ng/ml (40xMIC) was estimated by plating serial 10-fold dilutions of inoculum on drug-free and Q203-containing 7H11 agar plates. The plates were incubated at 32° C and CFU were counted after 10 weeks to determine the proportion of resistant CFU growing on drug-containing plates. For mutation analysis, DNA was extracted by boiling a few colonies in 100 μl of 1X TE buffer. Five μl of this was used for amplification by polymerase chain reaction (PCR). The entire *qcrB* gene along with 150 bp flanking region was amplified using the specific primers (Table S3). The resultant 2000 bp product was sequenced to identify the mutations in the gene.

### Statistical analysis

GraphPad Prism 6 was used for statistical analysis. When comparing mean CFU counts between three or more groups within the same mouse strain, one-way analysis of variance was used with Bonferroni’s post-test to adjust for multiple comparisons. When data were not normally distributed, as with relapse time points at which some mice had zero CFU and others had rebounding CFU counts creating a bimodal distribution, group CFU counts were compared using the non-parametric Kruskal-Wallis test. Similarly, an unpaired t-test or Mann-Whitney test was used as the parametric or non-parametric test when comparing CFU counts between two groups. Two-way analysis of variance was used with Bonferroni’s post-test to test for interactions between treatment regimen and mouse strain.

## Acknowledgments

This study was supported by the National Institutes of Health (R01-AI113266). O.K. was supported by a personal grant from the Bernhard-Nocht-Institute for Tropical Medicine, Hamburg, Germany. We gratefully thank the TB Alliance for providing Q203.

